# A cholesterol switch controls phospholipid scrambling by G protein-coupled receptors

**DOI:** 10.1101/2023.11.24.568580

**Authors:** Indu Menon, Taras Sych, Yeeun Son, Takefumi Morizumi, Joon Lee, Oliver P. Ernst, George Khelashvili, Erdinc Sezgin, Joshua Levitz, Anant K. Menon

## Abstract

Class A G protein-coupled receptors (GPCRs), a superfamily of cell membrane signaling receptors, moonlight as constitutively active phospholipid scramblases. The plasma membrane of metazoan cells is replete with GPCRs, yet has a strong resting trans-bilayer phospholipid asymmetry, with the signaling lipid phosphatidylserine confined to the cytoplasmic leaflet. To account for the persistence of this lipid asymmetry in the presence of GPCR scramblases, we hypothesized that GPCR-mediated lipid scrambling is regulated by cholesterol, a major constituent of the plasma membrane. We now present a technique whereby synthetic vesicles reconstituted with GPCRs can be supplemented with cholesterol to a level similar to that of the plasma membrane and show that the scramblase activity of two prototypical GPCRs, opsin and the β1-adrenergic receptor, is impaired upon cholesterol loading. Our data suggest that cholesterol acts as a switch, inhibiting scrambling above a receptor-specific threshold concentration to disable GPCR scramblases at the plasma membrane.

## INTRODUCTION

G protein-coupled receptors (GPCRs), a superfamily of cell membrane signaling receptors (1, 2), moonlight as phospholipid scramblases (3–5). When reconstituted into large unilamellar vesicles (LUVs), purified Class A GPCRs such as opsin, the β1- and β2-adrenergic receptors, and the adenosine A_2A_ receptor, facilitate rapid, bidirectional, trans-bilayer movement of phospholipids (4, 6–9). Molecular dynamics (MD) simulations of opsin suggest that a dynamically revealed groove between transmembrane helices 6 and 7 (TM6 and TM7) allows phospholipids to transit from one side of the bilayer to the other. This occurs according to the credit card model (3, 10, 11), with the headgroups of transiting phospholipids ensconced within the groove while their hydrophobic tails engage the membrane. GPCR-mediated lipid scrambling is constitutive, as studies of bovine rhodopsin show that scrambling is unaffected by bound retinal or light (6, 7). Consistent with this observation, MD simulations indicate that scrambling-associated conformational changes are distinct from those associated with G protein activation (3, 10).

The plasma membrane of all human cells is a strongly asymmetric phospholipid bilayer, with lipids like phosphatidylserine (PS) confined to the cytoplasmic leaflet (12–16). PS is exposed at the cell surface only on demand, in response to triggers such as apoptotic signals or Ca^2+^ influx which activate specific scramblases, members of the TMEM16 and Xkr protein families, while silencing lipid importers (flippases) which act to restore trans-bilayer lipid asymmetry (16–18). However, the ubiquitous expression of GPCRs in the plasma membrane - for example, HEK293T cells express ∼75 different types of GPCRs (19) - poses a dilemma: how does lipid asymmetry persist in a membrane which contains these constitutively active scramblases? It is possible that inward transport of phospholipids by flippases antagonizes the scramblase activity of GPCRs to maintain lipid asymmetry. Alternatively, the unique lipid composition of the plasma membrane, notably its high content of cholesterol (13, 20), may play a role. GPCR signaling is known to be affected by cholesterol, which can act directly by binding to the receptors and/or indirectly by modulating membrane physical properties (21–30). We therefore hypothesized that cholesterol inhibits the ability of GPCRs to scramble phospholipids at the plasma membrane. In support of this idea, our recent MD simulations of opsin suggest that cholesterol stabilizes the closed state of the phospholipid translocation pathway, thereby inhibiting lipid scrambling (31).

To test our hypothesis, we took a reductionist approach, asking whether cholesterol affects GPCR-mediated scrambling in LUVs. Individual vesicles in a sample reconstituted with more than one lipid component are known to be compositionally heterogeneous (32–34). Because this characteristic can affect the statistics of protein reconstitution (35, 36) and consequently interpretation of scramblase assay outcomes, we developed a technique to supplement the vesicles with cholesterol post-reconstitution, to a level similar to that found in the plasma membrane. We now show that the scramblase activity of opsin and the β1-adrenergic receptor (β1AR) is impaired when the vesicles are supplemented with cholesterol. Our data suggest that cholesterol acts as a switch, inhibiting scrambling above a receptor-specific threshold concentration to disable GPCR scramblases at the plasma membrane.

## RESULTS

### GPCR-mediated phospholipid scrambling in LUVs

We used a variation of a previously described one-pot protocol (Fig. 1A) to reconstitute opsin into LUVs containing a trace quantity of the fluorescent phosphatidylcholine (PC) reporter 1-palmitoyl-2-C6-NBD-PC (NBD-PC)(Fig. S1) (6–8) for scramblase activity assays. The principle of the scramblase assay is outlined in Fig. 1B. NBD-PC molecules in the outer leaflet of the vesicles are rapidly bleached by dithionite, a negatively charged membrane-impermeant chemical reductant (6, 37, 38). In protein-free LUVs where scrambling does not occur detectably on the timescale of the experiment, dithionite eliminates ∼50% of the fluorescence as NBD-PC molecules in the inner leaflet of the vesicles are protected (Fig. 1B, upper panel ’liposome’). As dithionite is added in considerable excess over NBD fluorophores, fluorescence loss follows pseudo first-order kinetics, with a time constant (tau)∼10-15 sec under our standard conditions. For LUVs reconstituted with GPCRs, all NBD-PC molecules are bleached, as those in the inner leaflet are scrambled to the outer leaflet (Fig. 1B, lower panel ’proteoliposome’). We previously reported that the rate of opsin-mediated scrambling is comparable to the rate of the dithionite-mediated bleaching reaction (6–8), allowing only a lower limit estimate of the scrambling rate (but see below). However, the scramblase assay also provides end-point data whereby fluorescence reduction in excess of that measured for protein-free liposomes is an indication of the fraction of vesicles reconstituted with a functional scramblase. Notably, this is a binary read-out at the level of individual vesicles as those with at least one scramblase register as fully active.

**Figure 1.**
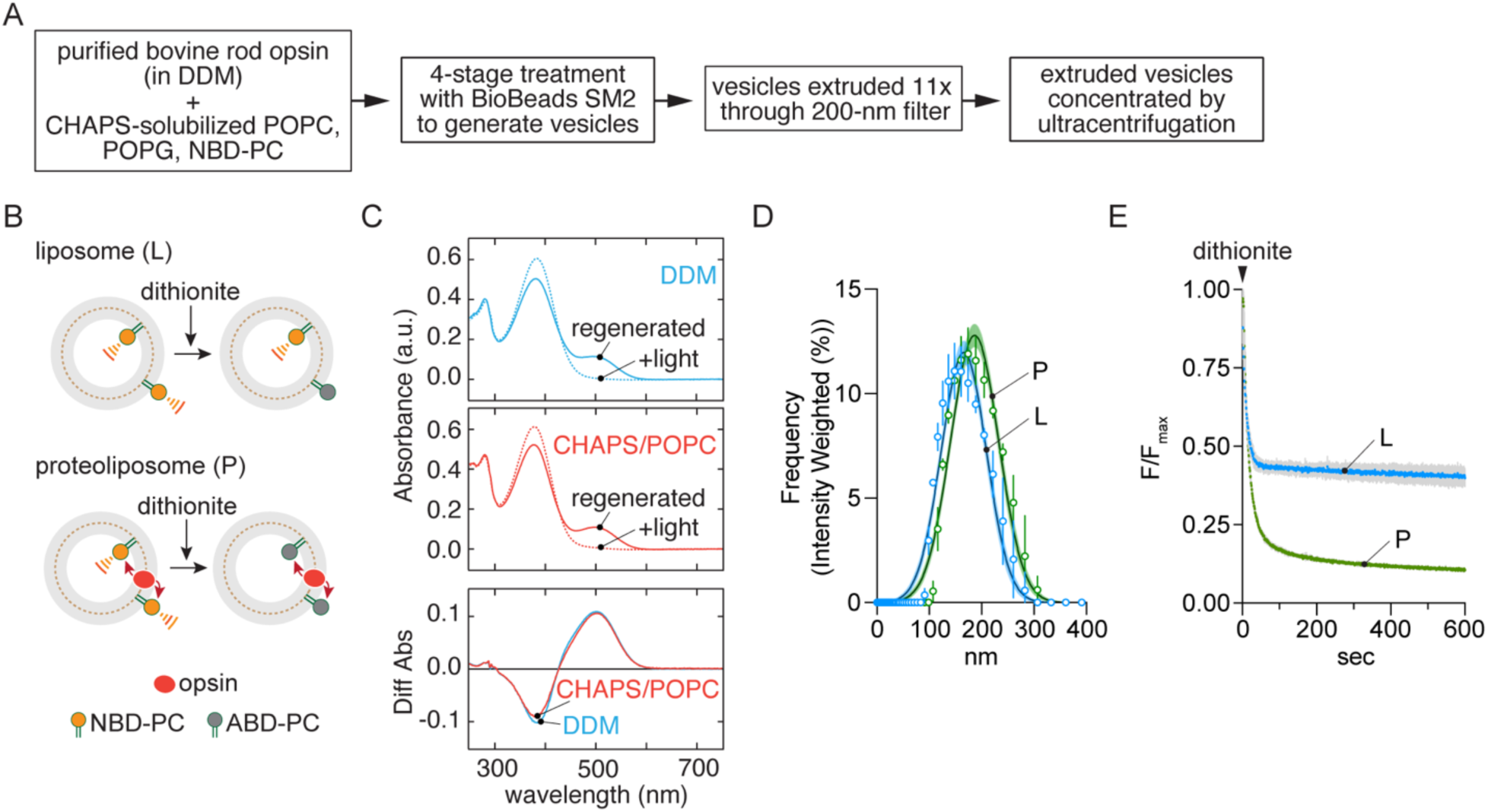
Opsin scrambling activity in reconstituted LUVs. **A.** Reconstitution protocol. **B.** Scramblase assay. Protein-free liposomes (L) or opsin-proteoliposomes (P) containing a trace quantity of fluorescent 1-palmitoyl-2-C6-NBD-PC are treated with dithionite which bleaches NBD-PC molecules in the outer leaflet of the vesicles to non-fluorescent ABD-PC. The extent of fluorescence loss is expected to be ∼50% for liposomes as dithionite cannot access NBD-PC molecules in the inner leaflet during the time frame of the assay, and ∼100% for opsin-containing vesicles in which NBD-PC molecules are scrambled between leaflets. **C.** Stability of opsin under reconstitution conditions. Regeneration of rhodopsin from opsin with 11-*cis*-retinal in DDM, or CHAPS supplemented with POPC. 11-*cis*-retinal (20 µM) was added to opsin and UV/vis spectra were measured after 30 min incubation at room temperature (“regenerated”). The sample was then illuminated with yellow light (>500 nm) for 1.5 min, and a fresh spectrum was recorded (“+light”). Regeneration was performed in 0.1% w/v DDM, 20 mM Hepes, pH 7.5, 150 mM NaCl (*top panel*), or after incubating the sample in 1.8% w/v (∼30 mM) CHAPS + 1.1% w/v POPC at 4°C for 1 h (*middle panel*). Difference spectra (*bottom panel*) were calculated by subtracting the spectrum obtained after illumination from the corresponding spectrum before illumination. The amount of rhodopsin regenerated in CHAPS/POPC was >96% of that regenerated in DDM. **D.** Vesicle size determined by dynamic light scattering. The data are presented as intensity weighted histograms (n=3 technical replicates); the lines correspond to Gaussian least squares fits (error bars = 95% confidence interval), indicating mean diameters (± standard deviation) of 166 ± 43 and 187 ± 46 nm for the L and P vesicles, respectively. **E.** Scramblase assay traces for L and P vesicles. Fluorescence (λ_ex_=470 nm, λ_em_=530 nm) was recorded at a frequency of 1 Hz for ≥ 100 s to obtain a stable initial value (subsequently used to normalize the traces) before adding dithionite at t=0 sec and monitoring fluorescence decay. Colored lines = mean fluorescence; grey bars = standard deviation (n=3 independent reconstitutions). Fitting to a two-phase exponential decay model (constrained to an initial value of F/F_max_=1.0) yielded plateau values of 0.40 and 0.10 for the L and P traces.

We purified bovine opsin from an n-dodecyl-β-maltoside (DDM) extract of bovine rod outer segments using Con A Sepharose chromatography. The protein was added to CHAPS-solubilized phospholipids, and the mixture was treated with detergent-adsorbing BioBeads. The resulting liposomes were extruded through a 200-nm filter and subsequently concentrated by ultracentrifugation (Fig. 1A). Several aspects of this preparation were characterized as follows. Purified opsin in DDM could be quantitatively regenerated to rhodopsin on adding 11-*cis*-retinal, indicating that the protein is well-folded and functional (Fig. 1C, top panel). This was also the case for opsin that had been diluted into the CHAPS/phospholipid mixture for reconstitution (Fig. 1C, middle panel; a difference spectrum (bottom panel) indicates that DDM-solubilized and CHAPS/phospholipid-solubilized opsins are equally well regenerated). The LUVs had a mean diameter of ∼175 nm as measured by dynamic light scattering (Fig. 1D). Opsin-containing LUVs (proteoliposomes, P) had scramblase activity as expected (6, 7), with dithionite addition causing a deep (∼90%) reduction in NBD-PC fluorescence (Fig. 1E). In contrast, ∼60% of fluorescence was lost on adding dithionite to protein-free liposomes (L) which were prepared in parallel (Fig. 1E, we expect ∼53% fluorescence loss for LUVs of the size that we prepare (39), but occasionally observe greater reduction - as seen here - for reasons that are not entirely clear). The fraction of vesicles (f) containing a functional scramblase was determined as f=1 - P/L_i_ from the exponential fit-derived plateau values, P and L_i_, of fluorescence obtained after dithionite treatment of proteoliposomes and matched protein-free liposomes, respectively. For the data shown in Fig. 1D, f=1-(0.1/0.4) = 0.75. Thus, 75% of the vesicles in the proteoliposome sample analyzed in this experiment possess a functional scramblase (7, 8). Our reconstitutions were set up to result in ∼15 copies of opsin per vesicle on average; however, as 25% of the vesicles lack scramblase activity, and presumably lack opsin, the average occupancy of the proteoliposomes is ∼20 copies of opsin/vesicle.

The fluorescence trace for opsin-proteoliposomes (Fig. 1E) could not be modeled using a 1- phase exponential function as previously described (6–8, 40), instead requiring a double-exponential fit with a slow ’scrambling’ phase characterized by a time constant, tau(slow), ∼150 s. The reason for this difference is unclear but could be due to the specifics of the updated reconstitution protocol and the nature of the opsin preparation (native bovine opsin versus HEK cell-expressed thermostable opsin used previously). Regardless, obtaining a measure of the scrambling rate is an advantage for our analyses as will become clear later.

### Methodology for testing the effect of cholesterol on scrambling

To test the effect of cholesterol on scrambling it is necessary to consider that LUVs assembled from two or more lipid components are compositionally heterogeneous (32–34). We assessed this heterogeneity by using total internal reflection fluorescence (TIRF) microscopy to image individual vesicles (Fig. 2A) in a preparation of cholesterol-containing LUVs with trace amounts of two fluorescent membrane dyes, Cy5-PE and TF-Chol (Fig. S1). The LUVs were captured onto a passivated glass slide (Fig. 2A) and imaged by TIRF illumination. We found a wide range of TF-Chol/Cy5-PE intensity ratios on quantifying individual vesicles that we localized as diffraction-limited spots (Fig. 2B,C). In addition, we found that the ratio of the two dyes was only weakly correlated (Fig. 2D,E). This evident dispersity in vesicle composition poses a technical problem for our study of the effect of cholesterol on scrambling. As protein reconstitution likely depends on the specific lipid composition of individual vesicles, vesicle occupancy statistics (35, 36) will be different in LUVs prepared with cholesterol compared with control LUVs. This makes it difficult to determine whether an observed cholesterol-dependent reduction in the fraction of scramblase-active vesicles is due to the effect of cholesterol on activity versus its effect on protein reconstitution. To circumvent this problem, we opted to reconstitute opsin first, using the standard protocol described above (Fig. 1A), and then introduce cholesterol by controlled, stepwise treatment of the LUVs with MβCD-cholesterol complex (CDC) (41, 42).

**Figure 2.**
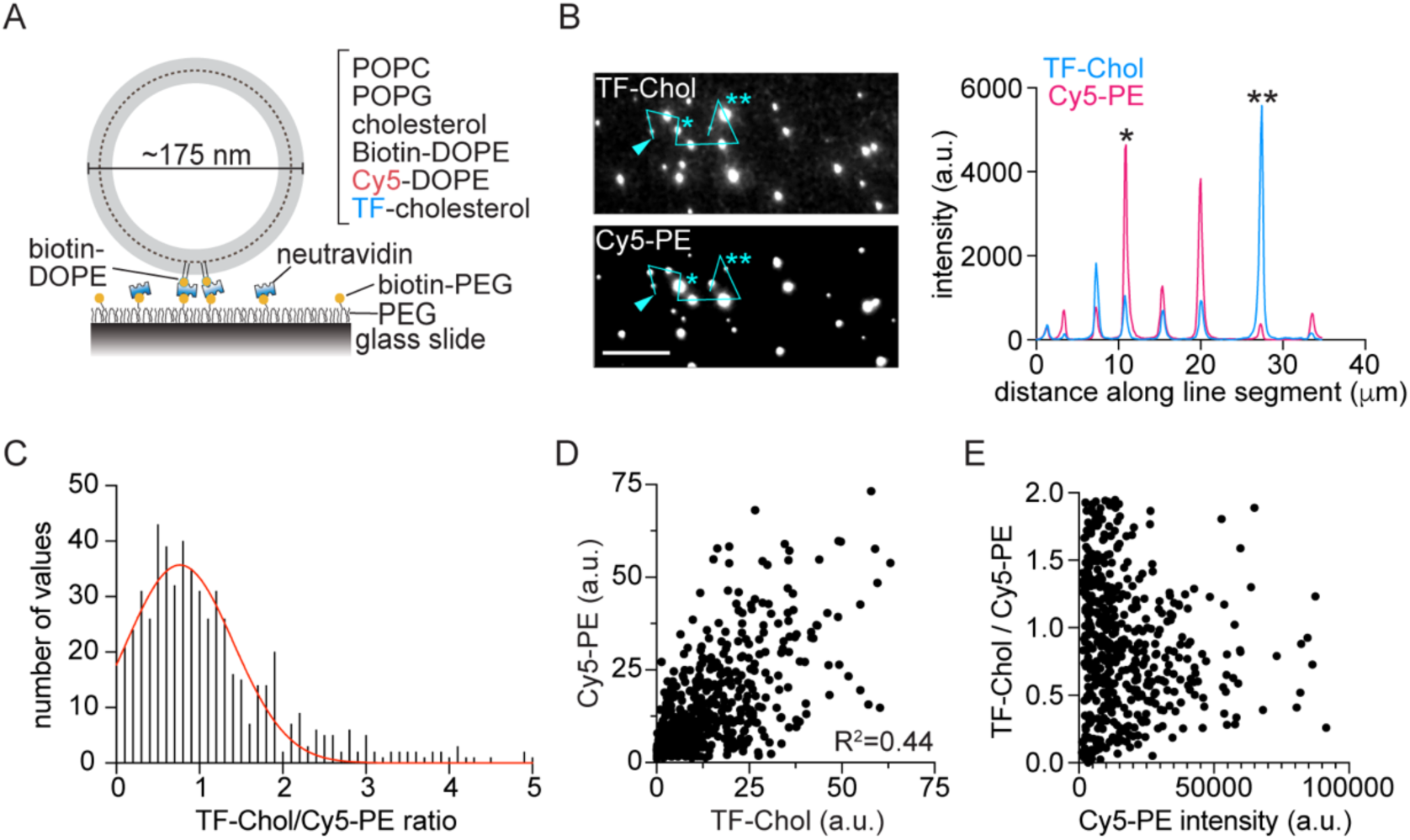
Compositional heterogeneity at the single vesicle level. **A.** Schematic showing capture of a single vesicle for TIRF imaging. POPC/POPG liposomes with 30% cholesterol and trace amounts of biotin-DOPE, Cy5-DOPE (Cy5-PE) and TopFluor cholesterol (TF-Chol) (see Fig. S1) were immobilized via neutravidin onto a passivated glass slide coated with a mixture of PEG and biotin-PEG. **B.** Representative fluorescence images of vesicles showing Cy5-PE and TF-Chol fluorescence. The images were analyzed using Fiji software. A line segment was drawn through the indicated vesicles (starting at the point indicated by the arrowhead) and the fluorescence intensity along the line was determined and graphed. Vesicles indicated by * and ** in the images are correspondingly indicated in the intensity profile. Scale bar 10 μ. **C.** Histogram of the TF-Chol/Cy5-PE ratio (each in arbitrary units), fitted to a Gaussian distribution (red line). The graph shows 576 of 595 vesicles that were imaged. **D.** Scatter plot of the TF-Chol and Cy5-PE intensities of individual vesicles. The correlation between the intensities of the two dyes in individual vesicles is low (R^2^=0.44). **E.** Scatter plot of the TF-Chol/Cy5-PE intensity ratio versus Cy5-PE intensity of individual vesicles, the latter providing a measure of vesicle size.

We established the cholesterol loading protocol using protein-free, NBD-PC-containing LUVs. A small volume of LUVs was rapidly mixed with freshly prepared CDC complex (∼0.4 mM cholesterol with an MβCD:cholesterol ratio = 10:1) and incubated at 30°C for 45 min, after which the vesicles were collected by ultracentrifugation. The supernatant was quantitatively removed before resuspending the vesicles in buffer. The mole percentage of cholesterol in the treated LUVs was determined using colorimetric assays for cholesterol (43) and phospholipid (44). A single incubation with CDC resulted in LUVs with ∼20% cholesterol; a second incubation increased the amount to ∼40%. LUVs mock-treated with buffer alone were processed in parallel. Cryoelectron microscopy (Fig. S2) imaging revealed that most (∼70-80%) of the vesicles are unilamellar, with a small fraction (∼20%) of bilamellar structures. The reason for the bilamellarity of a sub-population of the vesicles is not clear, but it is evident in mock-treated samples and therefore un-related to CDC treatment (Fig. S2).

Dithionite treatment (Fig. 3A) indicated that the fraction of NBD-PC in the inner leaflet (Lipid(inside), L_i_ (Fig. 3A, inset)) is similar in the initial and mock-treated LUVs, but higher in the CDC-treated sample (Fig. 3B), suggesting that the treatment results in loss of a fraction of NBD-PC molecules from the outer leaflet. The ratio of NBD-PC fluorescence to phospholipid content was the same in all samples (Fig. 3C), indicating that a similar fraction of phospholipids is also lost during CDC treatment. We confirmed this in experiments with *N*-NBD-PE, a fluorescent phospholipid which is expected to behave similarly to natural phospholipids in terms of its extractability (Fig. S3A). Thus, *N*-NBD-PE was extracted from the outer leaflet of LUVs on CDC treatment but not upon buffer (mock) treatment (Fig. S3A). We also tested the effect of treating LUVs with empty MβCD to mimic the effect of the high fraction of un-complexed MβCD that is present in CDC preparations. Similar to CDC-treated LUVs (Fig. 3A,B), MβCD-treated LUVs lost a fraction of their outer leaflet NBD-PC (Fig. S3B), resulting in a higher L_i_ value.

**Figure 3.**
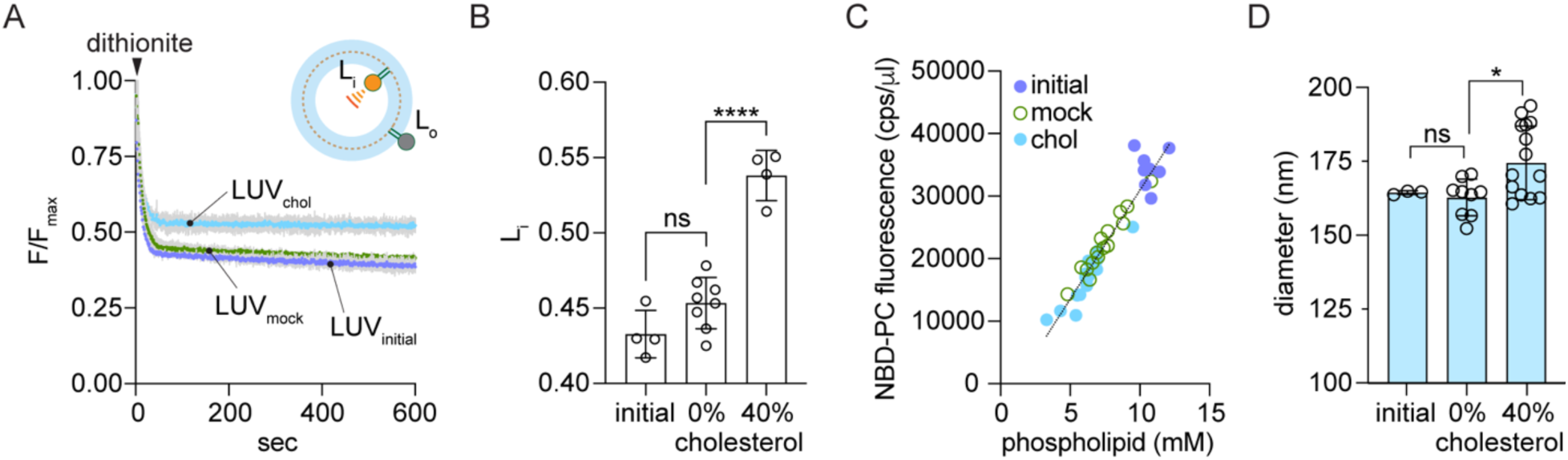
Methodology for cholesterol supplementation of LUVs. **A.** Protein-free liposomes were prepared by BioBead treatment of a CHAPS-solubilized mixture of bulk phospholipids and NBD-PC, followed by extrusion through a 200 nm filter. The LUVs (LUV_initial_) were subjected to two rounds of CDC treatment, or mock-treated with buffer, to generate LUV_chol_ and LUV_mock_, containing ∼40% and 0% cholesterol, respectively. The fraction of dithionite-accessible NBD-PC in the LUVs was determined by adding dithionite to continuously stirred LUVs in a fluorescence spectrometer and monitoring loss of fluorescence over time. Colored lines = mean fluorescence; grey bars = standard deviation (n=2 independent reconstitutions). Inset: Dithionite bleaches NBD-PC molecules in the outer leaflet (fraction L_o_) to non-fluorescent ABD-PC, leaving NBD-PC molecules in the inner leaflet unaffected (fraction L_i_; L_i_+L_o_=1). Symbols as in Fig. 1B. **B.** Fluorescence decay traces (panel A) were fit to an exponential decay model (single exponential decay plus a linear component with slope <10^-4^ s^-1^) and the plateau value (L_i_) was obtained from the fit. Data (mean ± S.D.) were obtained from 4 independent experiments (ns, not significant, ****, p<0.0001, ordinary one-way ANOVA using Tukey’s multiple comparisons test). **C.** Compiled data from four independent experiments. Liposome and proteoliposome samples were mock-treated or CDC-treated to load cholesterol, and the average fluorescence intensity (counts per sec, cps) (λ_ex_=470 nm, λ_em_=530 nm) averaged over ≥ 100 s was determined. The phospholipid concentration of the samples was determined using a colorimetric assay. The graph shows NBD fluorescence (cps per μl of vesicles used for measurement) versus phospholipid concentration (mM) for unprocessed (initial) vesicles, and mock-treated and cholesterol-loaded vesicles. The line indicates a linear regression fit of all the data. **D.** Liposome diameter determined by dynamic light scattering. The initial preparation of liposomes was compared with mock-treated or cholesterol-loaded samples (0 and ∼40 mole % cholesterol, respectively). The data represent the mean ± S.D. of triplicate measurements from 1-4 independent experiments (ns, not significant, *, p=0.03, ordinary one-way ANOVA using Tukey’s multiple comparisons test).

These results are consistent with the properties of MβCD, which can exchange both cholesterol (41) and phospholipids (45, 46). It is likely that asymmetric phospholipid loss from the outer leaflet during CDC treatment is compensated by cholesterol deposition as the diameter of CDC-treated LUVs is slightly larger than that of mock-treated samples (Fig. 3D). Thus, we envisage that cholesterol transfers from CDC to the outer leaflet of LUVs, and also rapidly equilibrates with the inner leaflet due to its high spontaneous flip-flop rate (46, 47). For subsequent experiments, as described below, we chose to compare CDC-treated proteoliposomes with correspondingly treated liposomes, the latter providing the means to calculate the fraction of scramblase-active vesicles in the proteoliposome sample as described above. We also prepared buffer-treated control samples of liposomes and proteoliposomes to obtain the fraction of scramblase-active vesicles in a compositionally non-perturbed sample which had otherwise been subjected to the same sample work-up as the CDC-treated vesicles. We chose not to use MβCD-treatment to generate control samples as this would result in unilateral lipid loss, unlike CDC-treatment where cholesterol deposition compensates for phospholipid loss.

### Cholesterol, above a threshold concentration, inhibits GPCR-mediated scrambling

We incubated opsin-containing and protein-free LUVs with CDC or buffer, using two rounds of incubation to reach ∼40% cholesterol in the CDC-treated samples. Mock-treated proteoliposomes had scramblase activity as expected (Fig. 4A) with a large fraction of the vesicles (∼0.87, calculated as described above) containing a functional scramblase (Fig. 4B). However, cholesterol supplementation lowered this value to ∼0.66 (Fig. 4A,B), indicating that ∼25% of the vesicles had been inactivated. There was no significant difference in the kinetics (tau(slow)) of the slow phase of dithionite-mediated fluorescence decay between the mock-treated and CDC-treated proteoliposomes, indicating that the CDC-treated vesicles that retained scramblase function are fully active (Fig. S4A,B). As the same reconstituted preparations were used for both mock-or CDC-treatment, this result can be parsimoniously explained by suggesting that cholesterol is heterogeneously incorporated into the vesicles, such that opsin-mediated scrambling is inhibited in vesicles with a cholesterol concentration above a certain threshold concentration. Thus, the two rounds of CDC-treatment used here result in ∼25% of the vesicles having a high, scramblase-inactivating cholesterol concentration.

**Figure 4.**
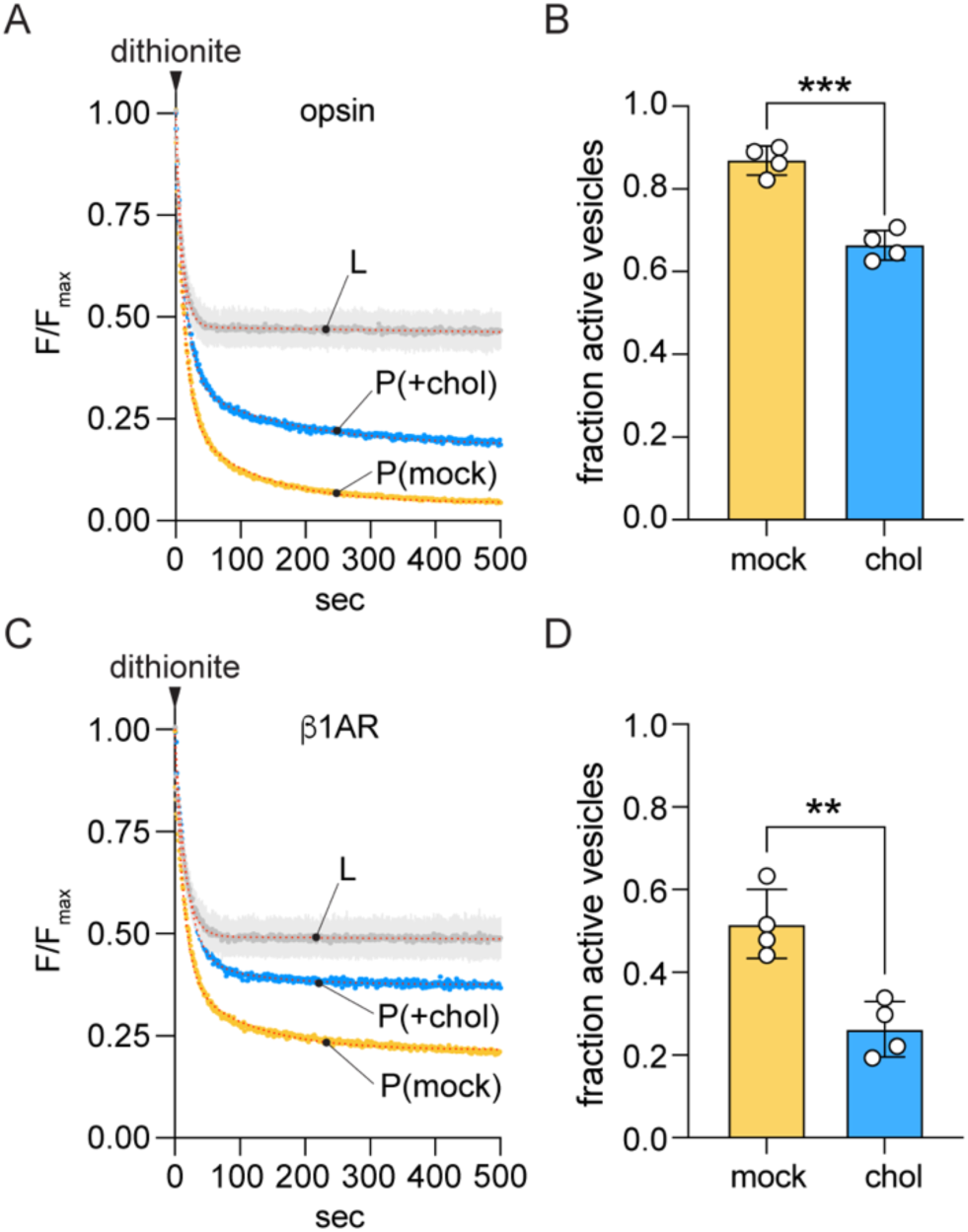
Cholesterol inhibits phospholipid scrambling by GPCRs. **A.** Representative fluorescence traces obtained on dithionite addition to liposomes (L) and mock-treated or cholesterol-loaded opsin-proteoliposomes (P(mock) and P(+chol), respectively). The cholesterol-loaded sample had ∼40% cholesterol. Data were obtained as described in Fig. 1D. The liposome trace is shown as the average (± S.D.) of the traces obtained for mock and CDC-treated liposomes for simplified presentation (see Fig. 3A for examples of individual traces). Dotted lines superimposed on the recorded data for P(mock) and P(+chol) represent double-exponential fits. The liposome data were fitted as in Fig. 3B. **B.** Bar chart of the fraction of active vesicles in P(mock) and P(+chol) samples (***, p=0.0002, unpaired t-test (two-tail)), determined as f=1 - P/L_i_ from the exponential fit-derived plateau values, P and L_i_, of fluorescence obtained after dithionite treatment of proteoliposomes and matched protein-free liposomes, respectively. **C.** As in panel A, except that β1AR-proteoliposomes were tested. **D.** As in panel B, except that β1AR-proteoliposomes were tested (**, p=0.0031, unpaired t-test (two-tail)).

As the effect of cholesterol on GPCR properties may be temperature dependent (48), we performed scramblase assays on mock-treated and CDC-treated opsin proteoliposomes at 15°C, 25°C and 35°C. The measurements (Fig. S5) show that scrambling is similarly inhibited by cholesterol at all three temperatures, resulting in ∼25% of the vesicles being inactivated. The observation of cholesterol-mediated inhibition of scrambling at 35°C, highlights its physiological significance.

To generalize our results with opsin, we tested the effect of cholesterol on scrambling by turkey β1AR, a Class A GPCR with previously reported scramblase activity (6). The receptor was purified in the presence of the agonist isoproterenol (49) and reconstituted (as described for opsin in Fig. 1A) and assayed also in the presence of 1 mM isoproterenol. The traces shown in Fig. 4C (P(mock) trace) confirm that β1AR is a scramblase. The extent of fluorescence decay for the mock-treated sample (Fig. 4C (P(mock)) was not as great as seen for a comparable opsin sample (Fig. 4A, (P(mock)), indicating that β1AR is less efficiently reconstituted. This may be because the protein multimerizes as detergent is withdrawn during the reconstitution process (see related analyses of opsin in references (7, 8, 50)), resulting in fewer vesicles being populated. However, as seen for opsin, the fluorescence decay trace required a double-exponential fit, with the slow second phase providing a measure of the scrambling rate. As also observed for opsin, the fraction of scramblase-active β1AR-containing vesicles decreased after two rounds of CDC treatment (Fig. 4C), but interestingly, the effect was greater, with ∼50% of the vesicles being inactivated compared with ∼25% for opsin vesicles. There was no significant difference in the scrambling rate between the mock-treated and CDC-treated β1AR-proteoliposomes, indicating that the CDC-treated vesicles that retained scramblase function are fully active (Fig. S4C), as also noted for opsin (Fig. S4A,B). We conclude that β1AR-mediated scrambling is more sensitive to cholesterol than opsin-mediated scrambling as ∼50% (versus ∼25% for opsin) of the twice-CDC-treated vesicles have a high, scramblase-inactivating cholesterol concentration.

### Heterogeneity in cholesterol content of vesicles

A key element of our interpretation of the effect of cholesterol on GPCR-mediated scrambling is that the reconstituted vesicles are compositionally heterogeneous with respect to cholesterol. To investigate this point further, we used single particle profiling (SPP)(33) to assess the membrane fluidity of individual vesicles, a measure of their cholesterol content. For this, we doped the vesicle samples with Nile Red 12S (NR12S)(51), an environment sensitive ratiometric dye, and interrogated hundreds of single vesicles as they diffused through the diffraction-limited observation volume of a confocal microscope. We measured the fluorescence intensity of NR12S at two emission bands (wavelengths 490-560 nm and 650-700 nm) and calculated the corresponding Generalized Polarization (GP), an index ranging from +1 to -1. GP is inversely proportional to membrane fluidity, increasing roughly linearly with the cholesterol content of POPC LUVs up to ∼35% cholesterol (33). GP distributions for CDC-treated vesicles (L_C_ and P_C_) were shifted to higher values compared to those for mock-treated samples (L_M_ and P_M_) (Fig. 5A-C). Notably, the shift in GP (ΔGP) was somewhat greater for proteoliposomes (ΔGP_P_ = GP_PC_-GP_PM_ = 0.27) than for protein-free liposomes (ΔGP_L_ = GP_LC_-GP_LM_ = 0.22)), suggesting a slightly higher (upto ∼5%) cholesterol loading of the protein-containing sample, which falls within the error of our colorimetric measurements of the average cholesterol concentration. The reason for the higher GP value of cholesterol-loaded proteoliposomes is not entirely clear - as the vesicles contain only ∼20 copies of opsin on average, it is unlikely that the presence of protein directly affects the extent of cholesterol loading.

**Figure 5.**
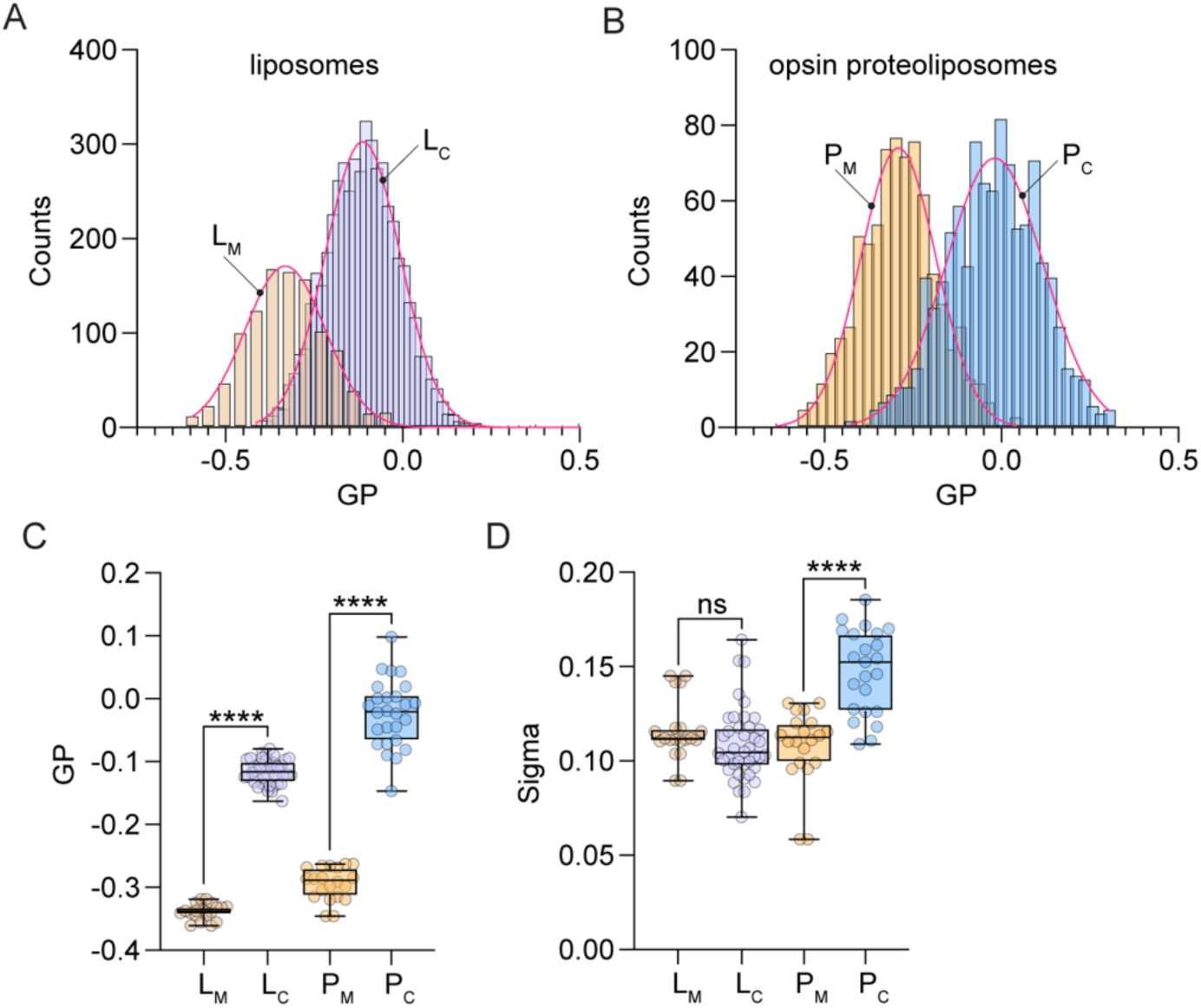
Heterogeneity of reconstituted vesicles. Mock-treated and CDC-treated liposomes (L_M_ and L_C_, respectively), and corresponding opsin-containing proteoliposomes (P_M_ and P_C_, respectively) were supplemented with 0.1 mol % of the ratiometric dye Nile Red 12S for measurement of Generalized Polarization (GP) by the Single Particle Profiling (SPP) technique. For panels C-D, Box = 25-75%, whiskers = min-max, line = mean value, statistical significance was probed by Kruskal-Wallis nonparametric test with multiple comparisons, **** corresponds to p<0.0001, ns for p>0.05. **A.** Histograms of GP for L_M_ and L_C_. **B.** Histograms of GP for P_M_ and P_C_. **C, D.** Mean GP and Sigma of GP, extracted from Gaussian fitting of histograms as in A and B after multiple technical replicates.

Multiple analyses generating replicate histograms, indicate a similar variation (Sigma) of the GP histogram for cholesterol-loaded and mock-treated liposomes (L_C_ vs L_M_, Fig. 5D), indicating that cholesterol-loading via the CDC method does not increase the pre-existing heterogeneity of the sample. However, a wider variation of the GP histogram was observed for the cholesterol-loaded versus mock-treated proteoliposome samples (P_C_ vs P_M_, Fig. 5D). The greater heterogeneity in GP of the cholesterol-loaded proteoliposomes may be due to the effect of protein on cholesterol loading (but see above), compounded by vesicle occupancy statistics where some vesicles do not have any protein, whereas the rest have a range of protein content.

The SPP results offer the following insights. First, GP measurements indicate significant compositional heterogeneity in the mock-treated samples, reflecting the heterogeneity of the starting reconstituted vesicle population. For perfectly homogenous one-component liposomes, Sigma is ∼0.07, compared with ∼0.12 for our mock samples (33). This is consistent with the imaging data of multi-component vesicles presented in Fig. 2. Second, as cholesterol would be expected to equilibrate between CDC and the vesicles, the heterogeneity in cholesterol content of the CDC-treated vesicles is likely due to the heterogeneity of the starting reconstituted vesicle population.

### Mechanism of cholesterol-mediated blockade of GPCR scrambling

Our data clearly indicate that cholesterol loading reduces the fraction of scrambling-competent proteoliposomes (Fig. 4). As the vesicles are compositionally heterogeneous (Fig. 5), these data appear to be consistent with a threshold mechanism whereby scrambling is blocked - on the time scale of our assay - in just those vesicles that contains a sufficiently high level of cholesterol. To test this idea, we contrasted it with an alternative mechanism whereby cholesterol has a graded effect such that the rate of scrambling is reduced gradually as cholesterol concentration increases. These two mechanistic scenarios can be distinguished in our scramblase assay as shown in Fig. S5. Thus, in the first scenario (threshold effect), the extent of fluorescence reduction in the scramblase assay (an indicator of the fraction of active vesicles) would be predicted to decrease as the average cholesterol content of the vesicle sample increases (Fig. S5A), but the scrambling rate would be unaffected (Fig. S5C). In the second scenario (graded effect), tau(slow) would increase in response to increased average cholesterol (Fig. S5C), but the extent of fluorescence reduction would remain the same (Fig. S5B).

We examined these predictions by titrating the average amount of cholesterol in the vesicles and assaying scramblase activity. Exemplary scramblase activity traces for once- and twice-CDC-treated opsin and β1AR vesicles are shown in Fig. 6A,B, with compiled results from several experiments presented in Fig. 6C,D. The data clearly show that the scrambling rate (reflected by tau(slow)) is unaffected by cholesterol (Fig. 6D), whereas the fraction of scramblase-active vesicles decreases as cholesterol concentration increases (Fig. 6C), supporting the threshold model (Fig. 6E).

**Figure 6.**
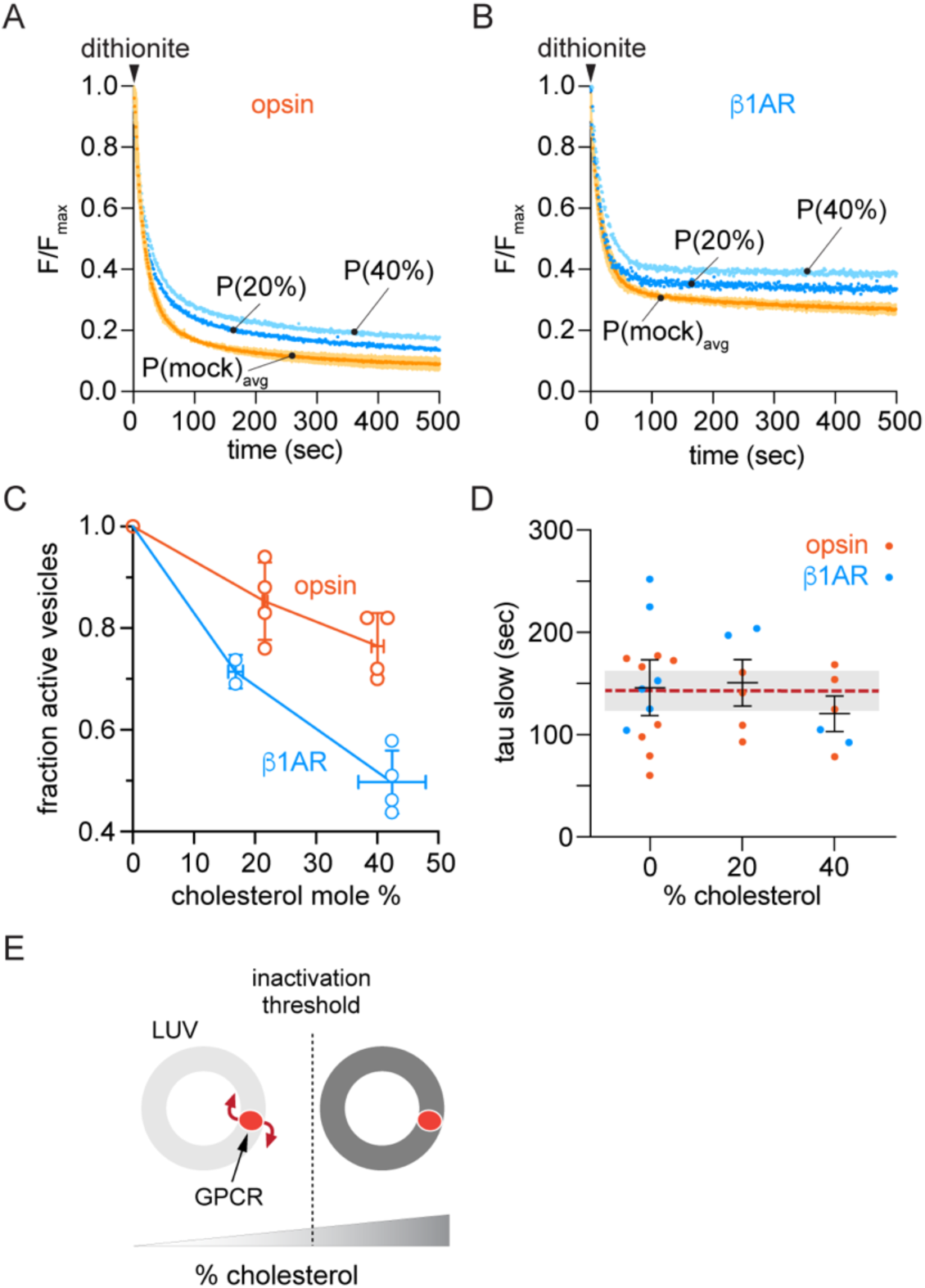
Cholesterol inhibits phospholipid scrambling by GPCRs in a switch-like manner. **A, B.** Representative fluorescence traces obtained on dithionite addition to mock-treated (P(mock)_avg_) or cholesterol-loaded opsin-proteoliposomes (A) or β1AR-proteoliposomes (B). The cholesterol-loaded samples, P(20%) and P(40%) had ∼20% or ∼40% cholesterol, respectively. Data were obtained as described in Fig. 1D. The mock-treated proteoliposome trace is the mean (± SD) of samples that were untreated or subjected to one or two rounds of mock-treatment. **C.** Cholesterol loading reduces the fraction of scramblase active vesicles. The ratio of the fraction of active vesicles in parallel preparations of cholesterol-loaded and mock-treated vesicles is graphed as a function of cholesterol concentration in the loaded vesicles (n=4 independent reconstitutions for all data points except n=2 for β1AR at ∼17% cholesterol). Mean values ± SD are shown for values along both axes. **D.** Fluorescence traces including those shown in panels A and B were analyzed by fitting to a double exponential decay function. The time constant (tau(slow)) associated with the slow phase of the exponential fit is shown as a function of the approximate average cholesterol concentration of the sample. The dashed line shows the average of all values, and the shaded bar indicates ± SD. **E.** Schematic illustration showing that GPCR scramblase activity is unaffected by cholesterol until a threshold level is reached, above which activity is blocked.

## DISCUSSION

We show that cholesterol acts as a switch, inhibiting GPCR-mediated scrambling above a threshold concentration. On incubating β1AR proteoliposomes once with CDC we obtained a sample with a mean cholesterol concentration of ∼20% in which scramblase activity was eliminated in ∼30% of the vesicles (Fig. 6C). Assuming that the range of cholesterol concentrations in individual vesicles within the sample follows a normal distribution, as seen in our GP measurements (Fig. 5B), it follows that the inactivation threshold concentration is >20%, reasonably estimated at ∼25% (Fig. S5D). This is lower than the average concentration of cholesterol in the plasma membrane which is ∼40% (20, 52–54). It is worth noting that local concentrations of cholesterol in the plasma membrane are even higher, for example in liquid ordered domains (55, 56) where GPCRs may be localized (57–59) or within a bilayer leaflet if cholesterol is not symmetrically distributed across the membrane (14, 20, 60), ensuring that GPCR scramblases are inactivated. We conclude that cholesterol-mediated inactivation of GPCR scrambling accounts for why GPCRs cannot dissipate the resting trans-bilayer lipid asymmetry of the plasma membrane.

Our results suggest several additional conclusions. First, it is unlikely that GPCRs scramble phospholipids within the endocytic system, a high cholesterol environment (61) similar to the plasma membrane. However, it remains possible that cholesterol-mediated inhibition of GPCR scramblase activity in the endocytic pathway may be relieved by one or a combination of factors, such as membrane curvature and/or the binding of transducer or accessory proteins that may stabilize scramblase-promoting GPCR conformations. Second, GPCR scramblases may play a role in cholesterol-poor membranes, such as those of the early secretory pathway. For example, newly synthesized GPCRs may contribute to the biogenic function of the endoplasmic reticulum, a membrane with just ∼5% cholesterol (7, 20, 62, 63), by scrambling phospholipids to promote membrane bilayer assembly (64–67). Third, the cholesterol concentration in bovine rod outer segment disks which house rhodopsin is low, ∼10% (68), suggesting that opsin-mediated phospholipid scrambling in disks occurs constitutively (69–71) to offset the bilayer disrupting activities of ATP-driven transporters (4, 72).

Although it is clear that cholesterol controls GPCR-mediated scrambling via a threshold effect, the titration data (Fig. 6C) suggest mechanistic complexities. Incubation of β1AR proteoliposomes twice with CDC results in an average cholesterol concentration of ∼40%, and a ∼50% loss of the scramblase-active population of vesicles (Fig. 6C). Assuming - as before - that the range of cholesterol concentrations in individual vesicles within this sample follows a normal distribution, we can readily see that the inactivation threshold concentration deduced from this sample is ∼40% (Fig. S5E), higher than the estimate of ∼25% that we obtained in samples treated once with CDC, while remaining below local cholesterol concentrations in the plasma membrane as noted above. This unexpected dose-dependent difference in our estimates of the inactivation threshold suggests that there may be more than one inhibitory mechanism at play. Cholesterol can influence the stability and quaternary structure of GPCRs by binding to the receptors and/or by modulating membrane physical properties (26, 73), and our MD simulations indicate that it can populate the proposed lipid transit groove between TM6 and TM7 in GPCRs to prevent scrambling (31). To understand how these mechanisms would result in different inhibitory thresholds, we suggest that it may be necessary to consider vesicle occupancy statistics (36, 74) which constitute another form of heterogeneity in our reconstituted system. Thus, we speculate that cholesterol-mediated changes in protein quaternary structure (73, 75–77) might inhibit scrambling at the lower threshold in vesicles with a ’high’ protein content, whereas for vesicles with fewer proteins, cholesterol - at a higher threshold - inhibits scrambling by populating the transit groove. Testing these ideas presents a significant challenge for future work, necessitating single vesicle approaches (74, 78) in place of ensemble methods. It is important to note that whereas we use cholesterol concentration as a convenient readout, it is the chemical potential of cholesterol that ultimately dictates its behavior in our reconstituted samples to drive the thresholds that we observe (79).

The inactivation threshold for opsin-mediated scrambling is clearly higher than that for β1AR, as only ∼25% of opsin-proteoliposomes are inactivated in samples with an average cholesterol concentration of ∼40% (Fig. 6C). What is the basis for this differential sensitivity to cholesterol? It is not unlikely that the two proteins differ in their mode of interaction with cholesterol as even the highly homologous β1- and β2-adrenergic receptors bind cholesterol in different ways (80). We note that we assayed β1AR in the presence of agonist to preserve its stability during purification/reconstitution, whereas opsin was processed ligand-free as a stable protein (Fig. 1C). Although we previously found that the scramblase activity of opsin is unaffected by the activation state of the receptor (6, 7), this aspect has not been systematically evaluated in other GPCRs and it is possible that ligand binding may affect their sensitivity to cholesterol. A further possibility is that β1AR and opsin may scramble lipids by distinct mechanisms. Sequence conservation analyses of Class A GPCRs (3, 10), reveal the presence of anionic glutamate at a membrane exposed location (residue 3.41 according to Ballesteros-Weinstein residue numbering) in the middle of the TM3 helix in β-adrenergic receptors; this residue is typically a hydrophobic or non-charged polar amino acid (*e.g.*, tryptophan in opsin) in other Class A GPCRs. Given its location and exposure to lipids, it is possible that the glutamate residue could attract water molecules to create the necessary hydrophilic environment for lipid translocation in β1AR, serving as alternative or additional to the TM6/7 pathway (10) (3) for lipid scrambling. This difference might contribute to the differential effect of cholesterol on β1AR-versus opsin-mediated scrambling.

In conclusion, we tested the hypothesis that cholesterol inhibits GPCR-mediated scrambling and identified that it does so via a threshold effect. Estimates of the inactivation threshold indicate that GPCR scramblases are effectively disabled in the cholesterol-rich environment of the plasma membrane. Our experiments highlight the methodological challenges in addressing questions such as this using ensemble methods. A detailed analysis of the molecular mechanism underlying the cholesterol effect(s) awaits future work.

## EXPERIMENTAL PROCEDURES

### Purification of opsin from bovine rod outer segment (ROS) membranes

ROS membranes were isolated from bovine retina by a standard discontinuous sucrose gradient method (81). The membranes were illuminated with yellow light (>500 nm) in the presence of GTP and washed to discharge transducin before treatment with 3M urea (pH 8.0) and repeated washing with low ionic strength buffer (pH 7.2) to remove residual membrane-associated proteins (82). The urea-stripped ROS membranes were incubated with 20 mM NH_2_OH in 20 mM HEPES pH 7.5, 150 mM NaCl, and again illuminated on ice to convert rhodopsin to opsin. Opsin was solubilized from the membranes with 1% w/v DDM, 20 mM Hepes pH7.5, 150 mM NaCl. After ultracentrifugation to remove insoluble material, the opsin-containing extract was applied to Con A Sepharose resin (previously equilibrated with 1% DDM buffer) by gravity flow. The column with bound opsin was washed with 20 bed volumes of buffer with 1% DDM, followed by 0.5% and 0.1% DDM in a stepwise manner to remove impurities including residual retinal oxime. Finally, opsin was eluted with buffer containing 0.3 M methyl-α-D-mannopyranoside and 0.1% DDM. The amount of intact opsin was assessed by the regeneration of rhodopsin with 11-*cis* retinal. 20 µM of 11-*cis* retinal was added to the opsin solution and UV/vis spectra were measured until the increase of 500 nm absorption reached the maximum. The results showed that the amount of rhodopsin regenerated in the mixture was 2.7 µM. Next, the effect of CHAPS/POPC mixture on regeneration was tested. The same amount of opsin was incubated for 1 h on ice in the presence of 1.8% CHAPS and 1.1% POPC, and then 20 µM of 11-cis retinal was mixed in. The regenerated rhodopsin was 2.6 µM in this condition, suggesting 96% of CHAPS/POPC-solubilized opsin could be regenerated.

### β1-adrenergic receptor (β1AR)

The turkey β1AR(H12) construct (PDB 7JJO) with an N-terminal T4-lysozyme fusion and C-terminal truncation (49), a gift of Drs. Minfei Su and Xin-yun Huang (Weill Cornell Medical College), was purified in the presence of 1 mM isoproterenol (Sigma 16504-500MG) after expression in insect cells as described (49).

### Reconstitution of opsin and β1AR into proteoliposomes

For a typical reconstitution experiment, two samples were prepared. For each sample, 432 μl POPC (25 mg/ml, Avanti Polar Lipids 850457), 48 μl POPG (25 mg/ml, Avanti Polar Lipids 840457), and 60 μl palmitoyl-NBD-PC (1 mg/ml, Avanti Polar Lipids 810130) were transferred to a 13 x 100 mm glass tube and dried under a stream of nitrogen for 30 min. The dried lipids were dissolved in 1 ml pentane and dried again for 30 mins, before adding 650 μl 70 mM CHAPS (Anatrace C316 5GM, ANAGRADE) prepared in Buffer A (50 mM Hepes, pH 7.4, 150 mM NaCl). The sample was shaken in an IKA Vibrax VXR agitator at 1500 rpm for 15 min at room temperature, before being sonicated in a bath sonicator (Elmasonic P30H, 37Hz, 100% power, pulse mode) for 5 min. The resulting clear solution was diluted to 10 mg/ml lipid and 35 mM CHAPS by adding 650 μl of Buffer A and transferred to a 2-ml eppendorf tube. Protein-free liposomes and opsin-proteoliposomes were generated by adding 100 μl of 0.1% DDM/PBS pH 7.2 or opsin (100 μl, 150 μg/ml in 0. 1% DDM/PBS pH 7.2), respectively. The samples were incubated for 45-60 min at room temperature with end-over-end mixing before adding BioBeads (Bio-Rad 152-3920) to remove detergent and generate vesicles. Prior to use, BioBeads were washed twice with methanol, twice with water and once with Buffer A (25 ml wash per g of BioBeads, gentle mixing for 15 min at room temperature per wash). BioBead addition was done in 4 stages at 4°C (2 h, 2 h, overnight (∼15 h) and 2 h), with end-over-end mixing, using 210 mg BioBeads per stage, and transferring the sample to 2-ml Eppendorf tubes with fresh beads each time. Following the final incubation with BioBeads, the now turbid samples were removed and extruded 11x through a 200 nm filter using an Avanti Mini-Extruder. The vesicles were concentrated by ultracentrifugation (TLA100.2 rotor, 75,000 rpm, 45 min, 4°C) and resuspension in 350 μl Buffer A. The resulting liposome and proteoliposome suspensions had a concentration of approximately 9-12 mM phospholipid, a PPR of ∼1.75 mg/mmol (∼15 opsin molecules per vesicle on average). Reconstitution of β1AR was done by the same procedure, with 1 mM isoproterenol included throughout, at a PPR of ∼2.7 mg/mmol (∼15 β1AR molecules per vesicle on average). The reconstitution is expected to be symmetric, with proteins inserted in either possible orientation as previously shown for opsin (7). As scramblase-catalyzed transbilayer movement is bidirectional, receptor orientation is not expected to affect the measurement of scramblase activity.

### Preparation of cyclodextrin-cholesterol complex (CDC)

A Hamilton syringe was used to transfer 250 μl of a 10 mM solution of cholesterol (Sigma C8667) in chloroform to a 13 x 100 mm glass tube. The solvent was evaporated under a stream of nitrogen before adding 5 ml of a 5 mM (6.55 mg/ml) solution of methyl-β-cyclodextrin (Sigma 332615, M_n_∼1310) prepared in Buffer A, and sealing the tube with parafilm wrap. The sample was sonicated in a bath sonicator (Elmasonic P30H, 37Hz, 100% power, pulse mode) for 15 min, until the dried cholesterol residue was finely dispersed, then placed in a rotary shaker, and incubated overnight at 35°C, 200 rpm. Following the overnight incubation, the tube was sonicated as previously for 5 min, before filtering the solution through an Acrodisc 0.2 μ syringe filter (Pall Corporation PN 4612). The resulting CDC preparation was used the same day. The cholesterol concentration in the preparation, measured as described below, was typically 0.3-0.4 mM.

### Cholesterol quantification

Cholesterol was quantified by the Zak colorimetric assay (43) with a linear response over a range of 0-100 nmol (0-39 μg). Briefly, CDC or vesicle samples (up to 100 μl aqueous volume) in 13×100 mm glass tubes were made up to 500 μl with absolute ethanol before adding 500 μl of the Zak reagent (prepared by dissolving 100 mg Iron(III) chloride hexahydrate (Sigma 44944-50G) in 4 ml of H_3_PO_4_ (Amresco E582-500ML) and 46 ml of concentrated sulfuric acid (Sigma 258105-500ML) in a glass-stoppered bottle and swirling to mix). A series of cholesterol standards in the range 0-30 μg (prepared from a 1 mg/ml solution of cholesterol in absolute ethanol) was assayed alongside. The samples were left at room temperature for 30 min before measuring their absorbance at 550 nm.

### Phospholipid quantification

Phospholipids were quantified by a colorimetric assay with a linear response over a range of 0-80 nmol (44). Samples were made up to 50 μl with water in 13×100 mm glass tubes, after which 300 μl perchloric acid (Sigma 30755-500ML) was added. The samples were heated for 1 h at 145°C. After incubation, 1 ml water was added, and the samples were allowed to cool to room temperature before adding 400 μl 12 mg/ml ammonium molybdate tetrahydrate (EMD Millipore AX1310-3) and 400 μl 50 mg/ml ascorbic acid (Sigma A5960-100G). After vortexing to mix, the samples were heated at 100°C for 10 min before measuring their absorbance at 797 nm. A series of standards (0-80 nmol inorganic phosphate) was prepared from 4 mM and 0.4 mM solutions of Na_2_HPO_4_.

### Loading vesicles with cholesterol

All treatments were done with matched sets of liposomes and proteoliposomes. Vesicles (100 μl) were rapidly pipetted into 1 ml of CDC in a 1.5-ml Eppendorf tube; a parallel sample was prepared using 1 ml Buffer A. The tubes were placed in an Eppendorf Thermomixer and agitated at 1000 rpm at 30°C for 45 min, after which the vesicles were collected by ultracentrifugation (TLA100.2 rotor, 75,000 rpm, 45 min, 4°C). After removing the supernatant, the vesicles were resuspended in 100 μl Buffer A using a mini-glass Dounce pestle. A single round of CDC treatment resulted in vesicles with ∼20 mole percentage cholesterol; an additional round of treatment was used to increase the amount of cholesterol to ∼40%. The mole percentage of cholesterol was calculated as 100•c/(c+p), where c and p are the concentrations of cholesterol and phospholipid, respectively, measured via the colorimetric assays described above.

### Scramblase assay

Vesicles (10-15 μl) were added to Buffer A (final volume 2 ml)) in a plastic fluorimeter cuvette equipped with a magnetic flea. The final concentration of vesicles was ∼100 μM phospholipid. For β1AR proteoliposomes and paired liposomes, 1 mM isoproterenol was included in the assay buffer. Time-based NBD fluorescence was monitored using a temperature-controlled Horiba Fluoromax fluorimeter equipped with an injection port. The following settings were used: λ_ex_ 470 nm, λ_em_ 530 nm, excitation and emission slits 2 nm, cuvette temperature 25°C, stirring rate 900 rpm, data acquisition frequency 1 Hz, recording. period 1000 s. After being allowed to stabilize for 2-3 min, fluorescence was recorded for ∼100 s before adding 40 μl of 1 M sodium hydrosulfite (Sigma 157953-5G) freshly prepared in 1 M Tris pH 10. All recordings were normalized to the average value recorded over the initial ∼100 s and analyzed in GraphPad Prism (version 9.1.0) by fitting to a one-phase exponential decay equation with a linear component (for liposome samples; the slope of the linear component was invariably weak (<10^-4^ s^-1^)) or a two-phase exponential decay equation (for proteoliposome samples), both constrained to a starting fluorescence value of 1.

#### Calculation of the fraction of active vesicles

The fraction of vesicles (f) containing a functional scramblase was determined as f=1 - P/Li from the exponential fit-derived plateau values, P and Li, of fluorescence obtained after dithionite treatment of proteoliposomes and matched protein-free liposomes, respectively.

#### Two-color single molecule TIRF imaging of cholesterols and liposomes

Cholesterol-containing LUVs were prepared with trace amounts of biotinyl-DOPE (biotin-PE)(Avanti Polar Lipids 870282), Cy5-DOPE (Cy5-PE)(Avanti Polar Lipids 810335), TopFluor-cholesterol (TF-chol)(Avanti Polar Lipids 810255) in the following molar proportions – POPC:POPG:cholesterol:biotin-PE:Cy5-PE:TF-chol (58.8 : 10 : 29: 1.0 : 0.2 : 1.0). Chloroform solutions of the lipids were combined in a 13 x 100 glass tube, and the solvent was evaporated under a stream of nitrogen. The lipids were then dissolved in pentane and re-dried under nitrogen. The resulting lipid film was resuspended in 50 mM Hepes, pH 7.5, 150 mM NaCl by shaking on a

IKA Vibrax VXR agitator at 1500 rpm, at room temperature for 1 h, then subjected to 5 freeze-thaw cycles, and extruded 11x through a 200-nm filter. For two-color TIRF microscopy, liposomes were immobilized on passivated glass coverslips prepared by coating with a mixture of biotinylated PEG and methoxyPEG via neutravidin as previously described (83). No liposomes adhered in the absence of neutravidin. After obtaining an optimal number of spots in the field of view, unbound liposomes were washed off with buffer (10 mM Tris, pH 7.5, 50 mM NaCl) prior to movie recording. Imaging was done using a 100x objective (NA 1.49) on an inverted microscope (Olympus IX83) in total internal reflection mode at 20 Hz with 50 ms exposure times with an sCMOS camera (Hamamatsu ORCA-Flash4v3.0). Liposome samples were excited with 640 nm and 488 nm lasers to excite Cy5 and TopFluor, respectively. Movies were recorded first with 640 nm laser and then with 488 nm laser to minimize potential FRET or direct excitation by the shorter wavelength laser. The intensity of TF-Chol and Cy5-PE in the collected images was analyzed using Fiji software (84). For each color, the background fluorescence intensity signal was subtracted from the movies. Next, a line segment was drawn through each liposome particle. The integrated intensity, which is the sum of the intensity values of the line, of individual liposomes was calculated for each frame within a movie. The maximum integrated intensity of each liposome within the movie was considered as the fluorescence intensity value.

#### Cryoelectron microscopy of liposomes

Quanifoil R 2/2 200 mesh copper grids (EMS #Q2100-CR2) were glow discharged at 25mA for 25s using a Pelco easiGlow system (Ted Pella). Liposomes at 5-7 mM were applied to the carbon side of a grid held in a Vitrobot Mark IV (ThermoFisher) chamber kept at 4°C and 100% relative humidity. Liposomes were applied three times (3 μl each time), with each application followed by a 1 min incubation and blotting. The sample was then immediately plunge frozen in liquid ethane. Samples were screened and imaged on a Talos Glacios 200kV microscope (ThermoFisher) equipped with a Selectris energy filter and Falcon 4i direct electron detector (ThermoFisher) at 63,000 X magnification, corresponding to 2A/pix.

#### Single Particle Profiling

Generalized polarization was quantified for single particles in solution using the Single Particle Profiler technique (SPP)(33). Briefly, vesicles were labeled with the environmental sensitive marker Nile Red 12S (NR12S). The fluorescence emission (λ_ex_ 488 nm) of the NR12S integrated into the LUVs was measured by letting the particles diffuse freely through the observation volume (focal volume of the 488 nm laser, roughly 200nm x 200nm x 500nm). The fluorescence intensity traces were recorded in two colour channels – 490-560 nm (green channel) and 650-700 nm (red channel). From intensity traces the individual peaks were isolated using home-made Python algorithm (freely available (33) ) and for each peak the generalized polarization was calculated using the expression GP = (I_green_ - I_red_)/(I_green_ + I_red_), where I_green_ and I_red_ are intensities in the green and red color channels, respectively. As each peak represents a single particle diffusing through the observation volume, a histogram of GP describes the population of all particles present in the solution. Furthermore, GP histograms were fitted with gaussian distributions so mean and sigma of these gaussians represented the GP value and heterogeneity of the LUV populations respectively.

## Data availability

All data are contained within the manuscript.

## Supporting information

This article contains supporting information (Figs. S1-S6).

## Acknowledgements

We thank Xin-yun Huang and Minfei Su (Weill Cornell Medical College) for 31AR, Laura Paweletz, Sarina Veit and Thomas Pomorski (Ruhr Universität Bochum) for the protocol to measure cholesterol via the Zak reaction, Andrey Klymchenko (University of Strasbourg) for the Nile Red 12S probe, Grace Dearden (Weill Cornell Medical College) for preparing Fig. S1, Milka Doktorova and Ilya Levental (University of Virginia School of Medicine), Sarah Veatch (University of Michigan) and Jeremy Dittman (Weill Cornell Medical College) for input, and Sam Canis for preliminary experiments. We acknowledge the support of the Cryo-EM Core Facility at Weill Cornell Medical College and thank Carl Fluck for assistance with Cryo-EM sample preparation, Glacios microscope operation and data collection, and Jared Shapiro (Weill Cornell Medical College) for quantifying the EM images. We thank the SciLifeLab Advanced Light Microscopy facility and Sweden National Microscopy Infrastructure (VR-RFI 2016-00968) for their support on imaging.

## Funding and additional information

We acknowledge support from National Institutes of Health grants R21 EY028314 (A.K.M., G.K.), R01 GM146011 (A.K.M.), the 1923 Fund (G.K.), a Rohr Family Research Scholar Award and the Irma T. Hirschl and Monique Weill-Caulier Award (J.L.), a Swedish Research Council Starting Grant (2020–02682), Human Frontier Science Program (RGP0025/2022), Karolinska Institutet and SciLifeLab (E.S.), and Natural Sciences and Engineering Research Council of Canada Discovery Grant RGPIN-2023-05764 and the Canadian Institutes of Health Research (CIHR) Operating Grant PJT-159464 (O.P.E.).

The content is solely the responsibility of the authors and does not necessarily represent the official views of the National Institutes of Health.

## Conflict of interest

The authors declare that they have no conflicts of interest with the contents of this article.

## Footnotes

### The abbreviations used are

ABD: 7-amino-2,1,3-benzoxadiazole
CDC: cyclodextrin cholesterol complex
DDM: n-dodecyl-3-maltoside
DPPE: 1,2-dipalmitoyl-sn-glycero-3-phosphoethanolamine
GP: Generalized polarization
GPCR: G protein-coupled receptor
LUV: large unilamellar vesicle
M3CD: methyl-3-cyclodextrin
NBD: 7-nitro-2,1,3-benzoxadiazole
PC: phosphatidylcholine
POPC: 1-palmitoyl-2-oleoyl-glycero-3-phosphocholine
POPG: 1-palmitoyl-2-oleoyl-sn-glycero-3-phospho-(1’-rac-glycerol)
ROS: rod outer segment
SPP: Single particle profiling
TF-Chol: TopFluor-cholesterol
TIRF: total internal reflection fluorescence

## Supplementary Information

**Figure S1.**
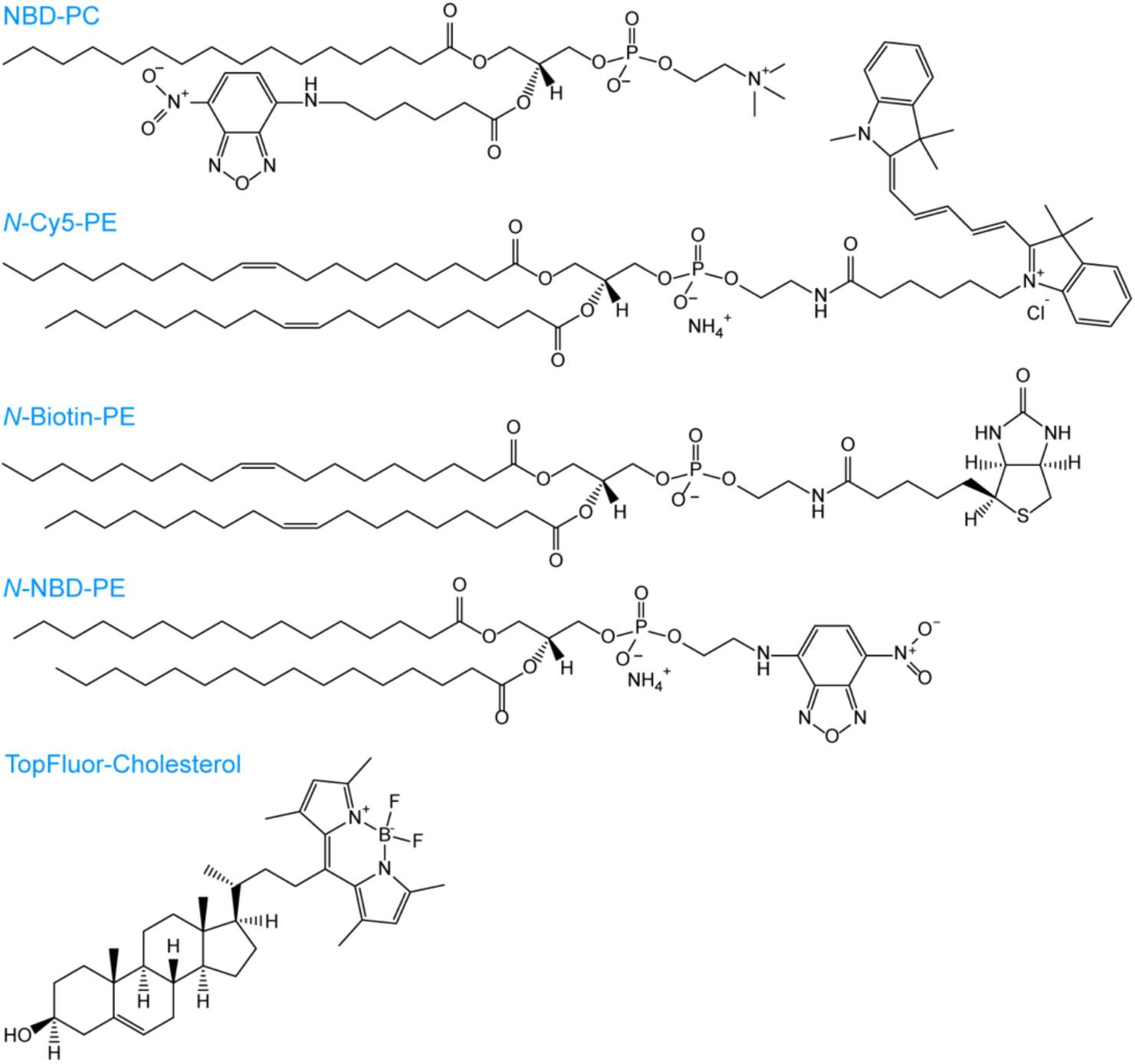
Fluorescent lipids used in this study. NBD-PC is the reporter lipid used for scramblase assays. *N*-Cy5-PE, *N*-Biotin-PE and TopFluor-Cholesterol are used in the single vesicle imaging experiment shown in Fig 2. *N*-NBD-DPPE is used in Fig. S3.

**Figure S2.**
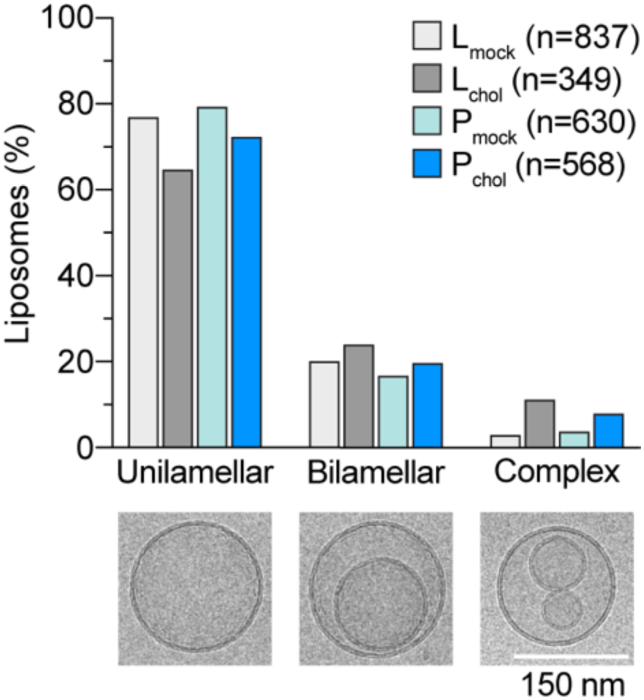
Morphology of mock-treated and CDC-treated vesicles determined by cryo-electron microscopy. Protein-free liposomes (L) and opsin-containing proteoliposomes (P) were subjected to two rounds of mock-treatment to generate L_mock_ and P_mock_ samples (0% cholesterol), or CDC-treatment to generate L_chol_ and P_chol_ samples (∼40% cholesterol). The samples were imaged by cryo-electron microscopy as described in Experimental Procedures. Examples of the different vesicle morphologies observed are shown in the images below the bar chart. The number of vesicles counted is indicated in the key.

**Figure S3.**
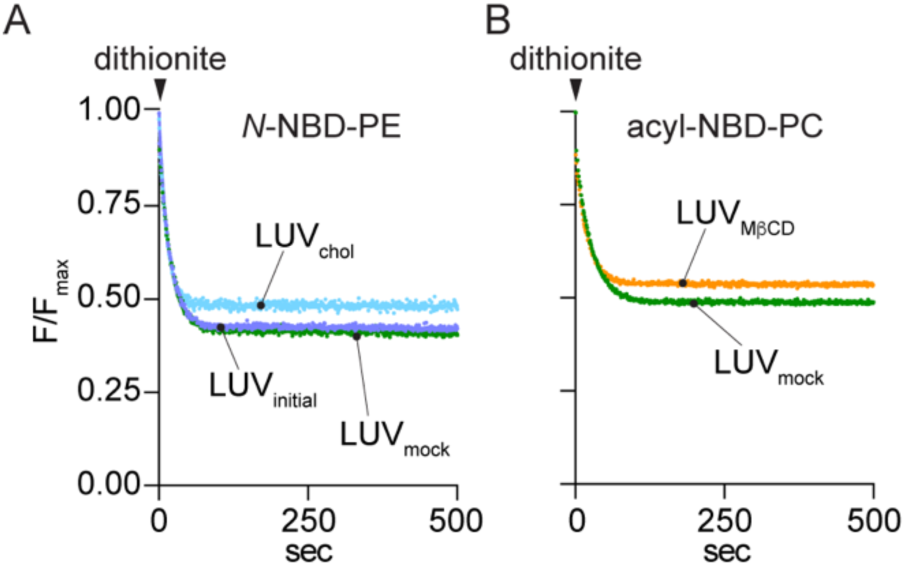
Controls for evaluation of the methodology for supplementing LUVs with cholesterol. **A.** LUVs containing *N*-NBD-PE were subjected to two rounds of treatment with buffer (LUV_mock_) or CDC complex (LUV_chol_). Dithionite was added and the time course of fluorescence decay was monitored. The data were analyzed as in Fig. 3A to determine L(inside)(L_i_, defined in Fig. 3A, inset)). Exemplary traces are shown. L_i_ = 0.43, 0.43 and 0.49 for LUV_initial_, LUV_mock_, and LUV_chol_, respectively. The relative ratio of NBD fluorescence to total phospholipid was 1.0, 0.99 and 0.98 for LUV_initial_, LUV_mock_, and LUV_chol_, respectively. **B.** LUVs containing acyl-NBD-PC were subjected to two rounds of incubation with buffer (LUV_mock_) or MβCD (LUV_MβCD_) and treated with dithionite as in Fig. 3A to determine L_i_. Exemplary traces are shown. L_i_ = 0.49 and 0.54 for LUV_mock_ and LUV_MβCD_, respectively.

**Figure S4.**
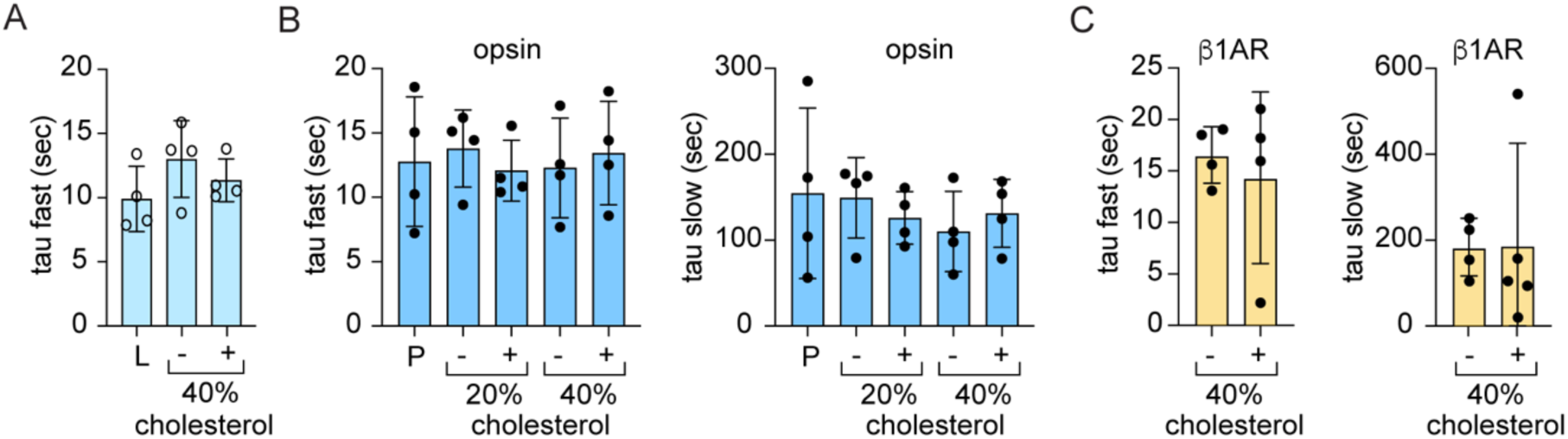
Kinetics of dithionite-mediated bleaching of NBD-PC in liposomes and GPCR-proteoliposomes. Protein-free liposomes (L) and GPCR-containing proteoliposomes (P) were subjected to one or two rounds of CDC-treatment to generate samples with ∼20% and ∼40% cholesterol, respectively. Mock-treated vesicles were prepared in parallel. Fluorescence traces including those shown in Figs 1E, 3A, 4A and 6D,E were analyzed by fitting to exponential decay functions. The liposome data (A) were fit to a mono-exponential function with a weak linear component (<10^-4^ s^-1^). The proteoliposome data (opsin (B) and β1AR (C)) were analyzed using a double exponential function. The time constants (tau values) associated with the fits are shown. Pairwise comparisons of tau values in each data set (ordinary one-way ANOVA (panels A,B), and two-tailed, unpaired t-test (panel C)) indicated no significant differences between values.

**Figure S5.**
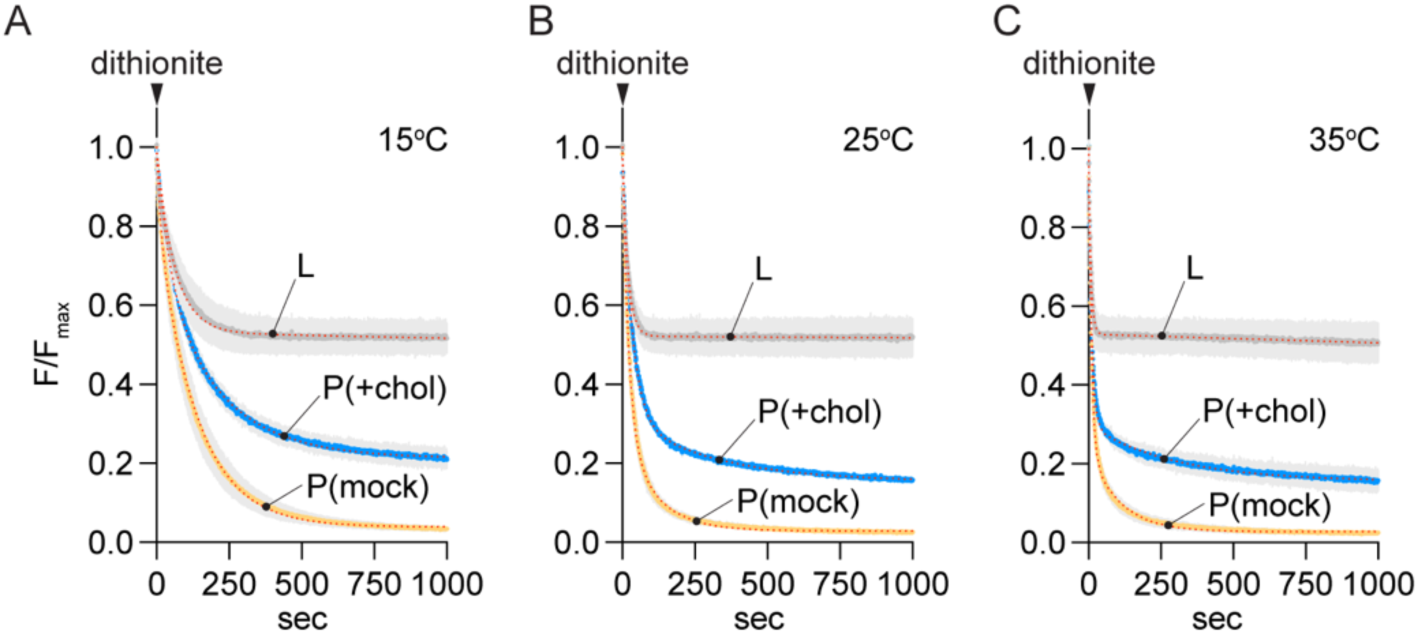
Cholesterol-mediated inhibition of opsin-mediated scrambling measured at different temperatures. Fluorescence traces obtained on adding dithionite to mock-treated or CDC-treated opsin-proteoliposomes (P(mock) and P(+chol), respectively) at different temperatures as indicated. The CDC-treated sample had ∼35% cholesterol. Data were obtained as described in Fig. 1D. Protein-free liposomes (L) were analyzed in parallel - for simplified presentation, as in Fig. 4, the liposome traces are shown as a single average (± S.D., error bars in light grey) of duplicate traces obtained for mock and CDC-treated liposomes. The P(mock) and P(+chol) traces are the average (± S.D., error bars in light grey) of duplicate measurements. The dotted line superimposed on the liposome data represents a mono-exponential decay model as in Fig. 3B. Dotted lines superimposed on the traces for P(mock) and P(+chol) represent double-exponential fits. The rate of dithionite-mediated bleaching of NBD fluorophores (obtained from fitting of the liposome data) increased with temperature: tau(fast) ∼45, 20, 5 s for measurements at 15°C, 25°C and 35°C, respectively. The tau(slow) values for the proteoliposome measurements were similar at all temperatures. The ratio of the fraction of active vesicles in P(+chol) versus P(mock) vesicles obtained from the three measurements was 0.74 ± 0.05 (mean ± S.D.)), consistent with the data shown in Fig. 6C.

**Figure S6.**
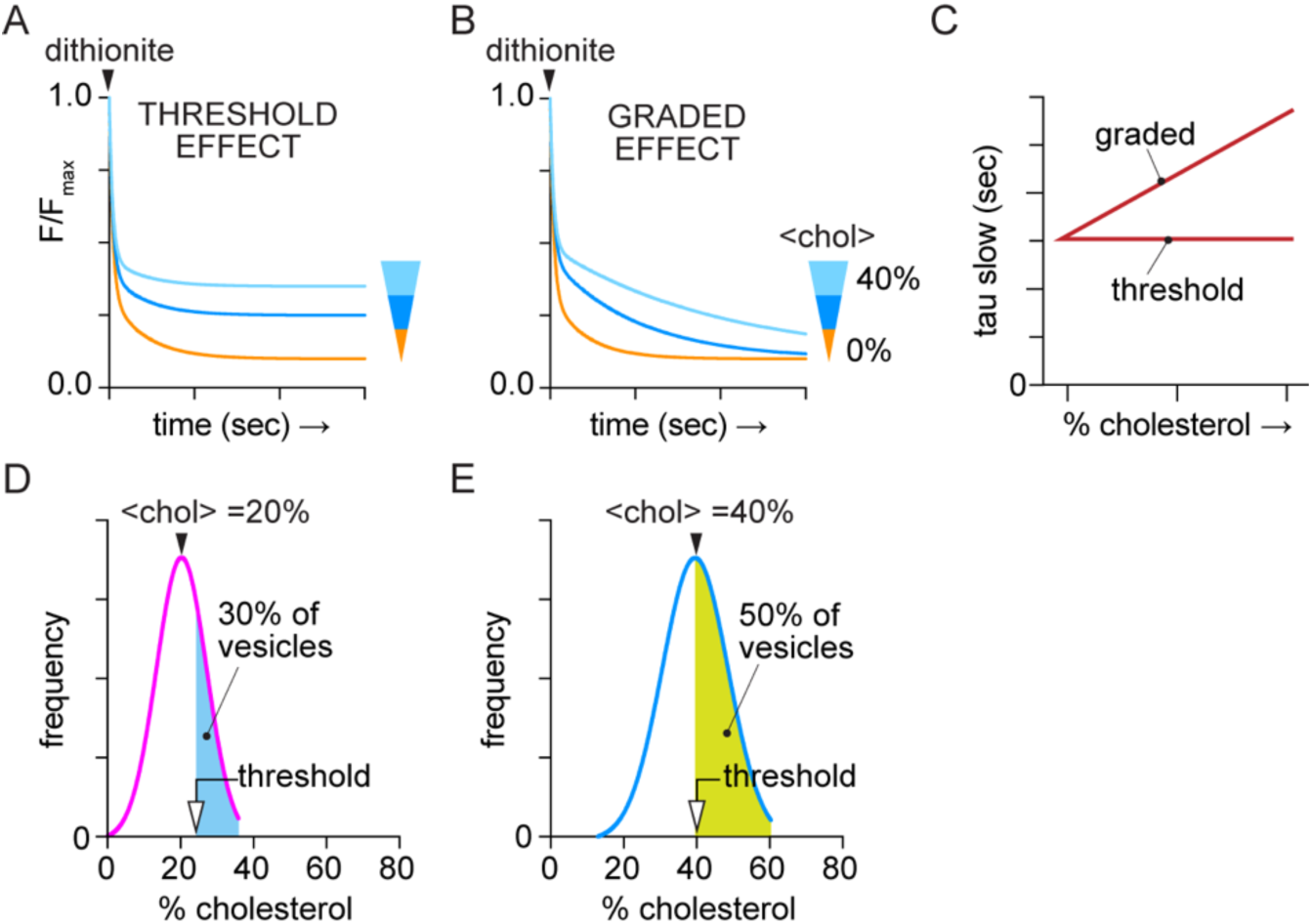
Predicted outcomes for threshold versus graded effect scenarios of cholesterol-mediated inhibition of GPCR scrambling. **A.** Threshold effect scenario where tau(slow) for a fraction of the cholesterol-treated vesicles is >100-fold greater than for mock-treated vesicles, hence not readily measured on the time scale of the assay. Treated vesicles that retain scramblase activity have the same tau(slow) as mock-treated vesicles. The schematic illustration shows a set of three traces, all with tau(fast)=20, tau(slow)=200, and plateau values of 0.1 (yellow), 0.25 (dark blue), 0.35 (light blue). **B.** A graded effect (panel A) would cause a gradual reduction in the rate of scrambling, seen as an increase in tau(slow) of a double-exponential fit (modeled here as a set of three traces, all with plateau =0.1 and tau(fast)=20, but with tau(slow)= 200 (yellow), 500 (dark blue), 1000 (light blue), in units of seconds. **C.** Predicted outcomes for tau(slow) as a function of cholesterol concentration according to the threshold or graded effect models. **D, E.** Models to estimate the inactivation threshold concentration of cholesterol for β1AR-mediated scrambling. The range of cholesterol concentrations in individual vesicles within the sample is presumed to follow a normal distribution. Panel D shows a schematic of a β1AR-proteoliposome sample treated once with CDC, resulting in a mean cholesterol concentration of 20%. The blue shaded area corresponds to 30% of the total area under the curve, representing the population of vesicles that is inactivated by cholesterol in this condition. Panel E shows a sample treated twice with CDC, resulting in a mean cholesterol concentration of 40%. The green shaded area corresponds to 50% of the total area under the curve, representing the population of vesicles that is inactivated by cholesterol in this condition. In both panels, the threshold concentration above which cholesterol inactivates scrambling in β1AR-proteoliposomes is indicated.

## Notes

### Competing Interest Statement

The authors have declared no competing interest.

### Summary of Updates

Supplemental Figure S5 has been added and thre are some minor text changes.

